# Trial by trial, machine learning approach identifies temporally discrete Aδ- and C-fibre mediated laser evoked potentials that predict pain behaviour in rats

**DOI:** 10.1101/2021.08.10.455801

**Authors:** A.C. Sales, A.J. Blockeel, J.R. Huxter, J.P. Dunham, R.A.R. Drake, A. Truini, A. Mouraux, RD. Treede, K.G. Phillips, A.E. Pickering

## Abstract

Laser evoked potentials (LEPs) – the EEG response to temporally-discrete thermal stimuli – are commonly used in experimental pain studies in humans. Such stimuli selectively activate nociceptors and produce EEG features which correlate with pain intensity. The rodent LEP has been proposed to be a translational biomarker of nociception and pain, however its validity has been questioned because of reported differences in the classes of nociceptive fibres mediating the response. Here we use a machine learning, trial by trial analysis approach on wavelet-denoised LEPs generated by stimulation of the plantar hindpaw of rats. The LEP amplitude was more strongly related to behavioural response than to laser stimulus energy. A simple decision tree classifier using LEP features was able to predict behavioural responses with 73% accuracy. An examination of the features used by the classifier showed that mutually exclusive short and long latency LEP peaks were clearly seen in single-trial data, yet were not evident in grand average data pooled from multiple trials. This bimodal distribution of LEP latencies was mirrored in the paw withdrawal latencies which were preceded and predicted by the LEP responses. The proportion of short latency events was increased after intradermal application of high dose capsaicin (to defunctionalise TRPV1 expressing nociceptors), suggesting they were mediated by Aδ-fibres (specifically AMH-I). These findings demonstrate that both C- and Aδ-fibres contribute to rodent LEPs and concomitant behavioural responses, providing a real-time assay of specific fibre function in conscious animals. Single-trial analysis approaches can improve the utility of LEPs as a translatable biomarker of pain.

## Introduction

Rapid heating of the skin by infrared lasers causes selective activation of thermally-responsive nociceptors. In humans this generates the percept of pain and triggers distinct EEG responses known as ‘laser evoked potentials’ (LEPs)^1,2^. Features of the LEP – particularly its amplitude – correlate with pain perception and this methodology has been employed in a diverse range of mechanistic pain studies^3–10^. In rodents, similar LEP features are reported to be closely related to nociceptive intensity^11–13^ and this similarity presents the possibility that the LEP could be used as a translatable biomarker of pain i.e. a proxy measure for pain applicable across species. In experiments focusing on pain mechanisms or novel analgesic drugs, the existence of a reliable, translatable marker could help bridge the gap between animal studies and clinical trials. To have confidence that the LEP is such a marker, it is necessary for it to have comparable underlying neural mechanisms - from the activation of nociceptors in the periphery, to the subsequent cortical event observed on the EEG. In humans, laser stimulation selectively activates Aδ- and C-nociceptors in superficial layers of the skin^2,14^, with EEG responses primarily reflecting the activation of Aδ fibres. LEPs on the timescale of C-fibres are generally only apparent in humans when A-fibres have been blocked, or when stimuli are carefully tailored to preferentially evoke C-fibre responses^1,15,16^.

In rodents, there is good evidence that contact heating of the paw can activate both C and Aδ-fibres (dependent on the rate of heating) to trigger withdrawal^17,18^. Laser stimulation of the plantar surface of the foot elicits responses in dorsal horn neurons at conduction velocities consistent with both Aδ- and C-fibres^19^. Other studies have reported evidence of C-fibre activation, but have described Aδ responses as highly variable, requiring higher intensity stimuli^19–21^. In addition, EEG and current sink studies have reported both short and long latency cortical responses, attributed to Aδ- and C-fibres respectively^22,23^. Challenging this view, two recent papers have proposed that the short latency LEP components in rodents are artefactual, created by the ultrasonic noise associated with laser stimulation^11,12^. Indeed this analysis has gone further to report that C-fibres are the sole mediators of the rodent LEP^11,12^. Clearly this would represent a substantial inter-species difference between human and rodent models.

The interpretation of EEG results is complicated by the common practice of averaging over events to form an overall LEP for a given stimulus energy. This process is designed to reduce the noise that is often present in EEG recordings, however it can also mask meaningful information. This is especially true when a small proportion of responses are qualitatively different from the rest. In this case, these rarer responses are liable to be ‘lost’ in the averaging, leading a reduction in information about response variability and potentially confounding interpretation of the resultant data^24,25^.

In this study, LEPs and behavioural responses were evaluated across a range of laser energies. Both individual and mean responses were analysed to gain insight into the relationships between stimulus energy, LEP morphology, behavioural responses, and the mode of transmission. We found mutually exclusive short and long latency responses occurring at the level of individual LEPs. Short latency events were rarer than long latency events (around 30% of events at higher stimulus energies) and were not initially apparent in the averaged data. Using a machine learning approach, we show that individual denoised LEPs can be used to predict behavioural responses with an accuracy of >70% - and that both the amplitude and latency of individual LEPs are important for classification. Injection of a high dose of capsaicin into the hindpaw – to defunctionalise TRPV1 expressing nociceptors – increased the proportion of short latency responses and caused the mean LEPs to become shifted towards shorter latencies. This provides evidence for the involvement of both Aδ- and C-fibres in rodent LEP responses and behaviour, and suggests that single trial analysis of LEPs can provide valuable information about the mode of transmission of nociceptive information from the periphery.

## Results

### Behavioural responses to laser stimuli

Infra-red Nd:YAP laser stimuli were delivered alternately to either the left or right hind paws at a range of energies between 0.75J and 2J (wavelength 1340nm x 4ms), with a minimum interstimulus interval of 30s (Figure 1a,b). Behavioural responses were initially scored into five categories, where a higher score corresponded to a greater degree of pain related behaviour (Figure 1c,d). However, because very few events were categorised as ‘1’ or ‘3’ (Figure 1d), these scores were too infrequent to allow meaningful statistical inference and so were ‘collapsed’ to produce a simplified 3-point scale (Figure 1c). In this simplified scale, a score of zero for a particular event indicated no response. Scores 1-2 denote a ‘flinch’ response, that is, a brief indication of awareness to the stimulus (for instance, a head turn, body movement or a momentary foot-lift), but without clear signs of pain-like behaviour. In contrast scores 3-4 describe a clear withdrawal of the foot with nocifensive responses such as licking, grooming, or extended attention to the affected foot (see Methods).

**Figure 1.**
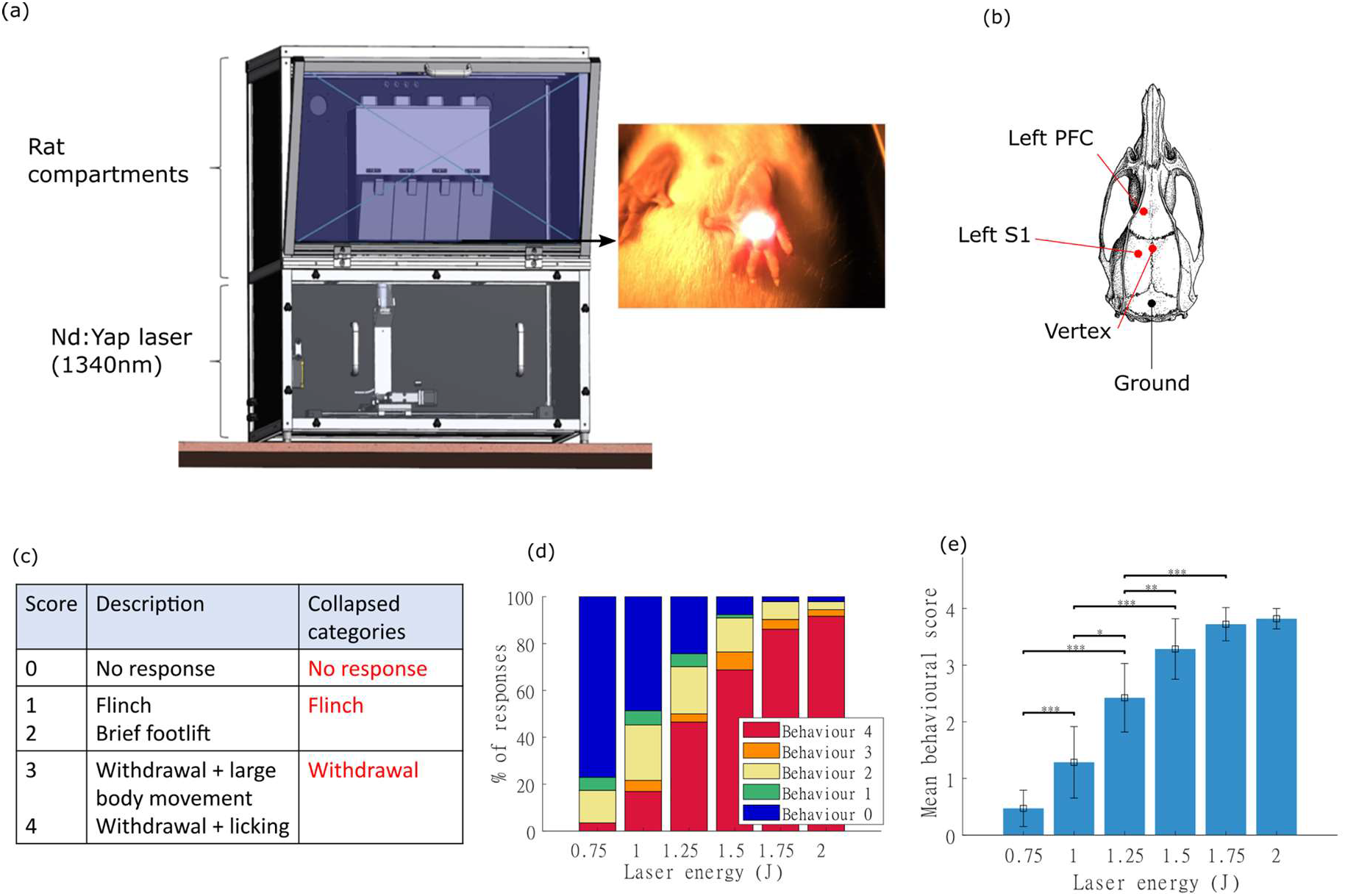
Overview of experiment and behavioural responses. (a) Apparatus overview. Four individual rat compartments were located on a glass floor above a fibre optic cable which transmitted the light from an infra-red Nd:YAP laser. With the aid of a camera in the lower compartment, a motorised xy-stage allowed remote targeting of the laser to the plantar surface. (b) Location of selected skull screws implanted in rats and used for recording EEG (c) Descriptions of behavioural score categories of the response to laser stimulation, with ‘full’ and ‘collapsed’ categories. (d) Proportions of behavioural responses in each category for each laser energy (across all animals) showing the increase in the behavioural response with laser energy. Note few responses were scored as ‘1’ or ‘3’. (e) Distribution of mean behavioural scores (repeated measures ANOVA, Bonferroni correction for multiple comparisons, * p<0.05, ** p<0.01, ***p<0.001)

The relationships between laser energy, side of stimulus and behavioural response were explored. As expected, there was a significant effect of laser stimulus energy on behavioural score (F=130.45, p<0.001, repeated measures ANOVA), with increasing laser energies evoking greater behavioural responses (Figure 1d,e). This stimulus-response function showed a sigmoid relationship: at the lowest stimulus energies, the behavioural responses were almost always ‘0’ (no response), with a steep rise between 1 and 1.5J, whilst at 1.75 or 2J responses plateaued and were almost always ‘4’ (withdrawal with licking). There was no significant effect of side of stimulation on the behavioural response score.

### Characteristics of laser event related potentials (LEPs)

To provide a comparison to previous studies^11–13^, the LEP responses for each animal were initially analysed by averaging EEG responses over all events within a given group - for instance, all events at a specific energy or behavioural category. Consistent with previous reports^11^, mean LEPs recorded across all skull sites had a stereotyped morphology that was comprised of a distinct peak whose amplitude and latency were related to the laser energy and behavioural response (Figure 2a-d). For the vertex EEG site, there was no significant lateralisation of response (as measured from peak LEP height, p>0.05, one way repeated measures ANOVA with laser energy and side as within subject factors), therefore for all subsequent analyses the vertex responses to both right and left hind paw stimulation were pooled. Lateralisation was found for left prefrontal and left somatosensory cortex LEP, with amplitudes that were greater following stimulation of the contralateral hind paw (F=6.1, p=0.03 and F=12.0, p=0.005 respectively, figure 2d). For these sites, all subsequent analyses were restricted to trials where the stimulation was applied on the contralateral side.

**Figure 2.**
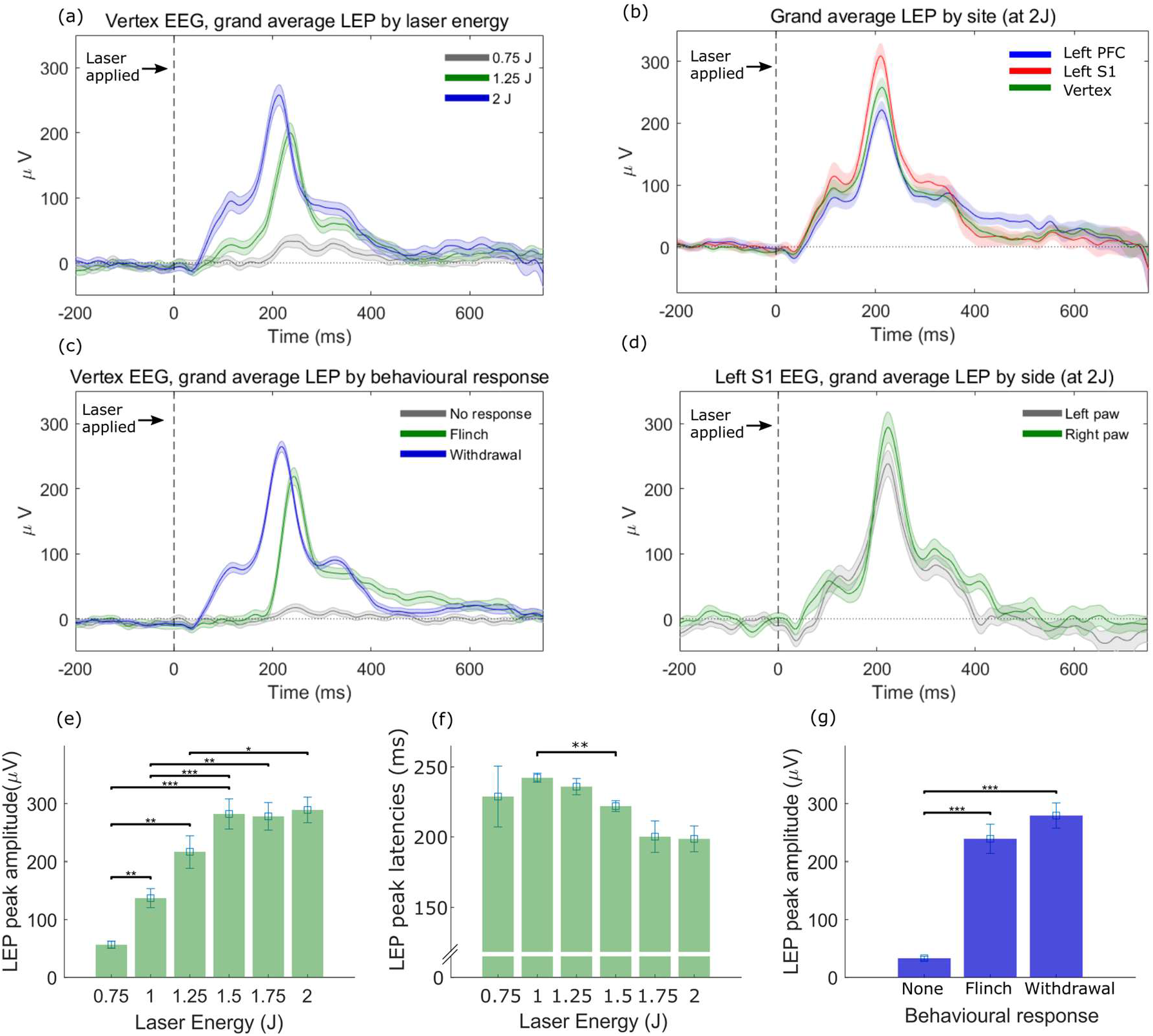
Grand average LEPs and variation of amplitude and latency. Form of the LEP waveform with respect to (a) laser energy, (b) EEG recording site, (c) behavioural responses and (d) side of stimulation ((a), (b) and (d) show responses at fixed laser energies, across all behaviours, while (c) shows responses associated with particular behaviours irrespective of laser energies). (e)–(g) Relationship between amplitude/latency of averaged vertex LEPs and laser energy or behaviour (repeated measures ANOVA, Bonferroni correction * p<0.05, ** p<0.01, ***p<0.001).

There was a positive correlation between laser stimulus energy and LEP amplitude (for the vertex site, F=32.8, p<0.001, repeated measures ANOVA, Figure 2e) in agreement with previous studies^11,13^. There was also a concomitant modest shortening in the latency to peak amplitude as the laser energy was increased (for the vertex site, F=2.6, p=0.04, Figure 2f).

### LEP amplitude is more strongly related to behavioural response than laser energy

The magnitude of the laser stimulus strongly influenced the behavioural response (Figure 1d,e), therefore we also expected there to be a correlation between LEP amplitude and behaviour. Indeed, when LEPs were grouped according to behaviour (averaging over intensities), there was a strong effect of behavioural category on LEP amplitude (for the vertex site, F=72.3, p<0.001, Figure 2g) but not on peak latency. Further analysis indicated that whilst both laser energy and behavioural response influence the amplitude of the LEP; behavioural response is the dominant factor. When laser energy was held constant within the range 1-1.75J (intensities at which all behavioural categories are observed and statistical analysis is meaningful), there was a significant interaction between behaviour and LEP amplitude (for fixed laser energies of either 1-1.25J: F=41.6, p<0.001, or 1.5-1.75J: F=31.8, p<0.001). In contrast, when the LEP data was analysed according to behavioural score there was no significant effect of laser energy on LEP amplitude, suggesting that LEP amplitude is more strongly related to the behavioural response. The relationship to laser energy is therefore secondary to the observation that higher energies are more likely to produce a stronger behavioural response (i.e. withdrawal) and thus a higher peak amplitude. Therefore as is believed to be the case in humans^26^, LEP characteristics likely reflect processing and decision making on incoming nociceptive information (reflected in behavioural response), rather than simply encoding nociceptive sensory information from the periphery.

Similar relationships between LEP amplitude, stimulus energy and behaviour were found for the left sensory and prefrontal EEG recording sites (supplementary table 1, with the exception of the lack of latency changes with stimulus energy within the sensory cortex). This suggests that representative LEP information can be usefully measured from a single recording site. Consequently, in the following sections the results will be based on data from the vertex EEG site unless otherwise indicated.

EEG spectral power analysis showed that laser stimuli were associated with a change in power in the delta (0.5-5Hz), theta (5-12Hz) and high gamma (50-100Hz) ranges, as previously reported^13^. The power in all ranges was increased relative to baseline in the 400ms time window following the laser stimulus (shown in supplementary figure 1). For the vertex site, the percentage increase in all ranges relative to a pre-stimulus baseline was related to both laser energy and behaviour (supplementary table 2a), however, the stimulus energy did not modulate power in the delta and theta ranges when analysed within behavioural response. In contrast, the relationship between spectral power and behavioural response remained significant for all three frequency bands when stimulus energy was held constant at 1J (at which a range of behaviours were evoked supplementary table 2b). This indicates that behaviour is the dominant factor determining the form of the LEP in both the amplitude and frequency domains.

### Machine learning investigation of LEP features that predict behaviour

We assessed whether any LEP features could reliably be used to classify the behavioural responses to individual events using machine learning algorithms, because such techniques can provide new insights, without pre-existing biomechanistic bias. Decision trees provide a straightforward method of achieving this aim and are importantly able to provide a clear account of the components of the LEP that are most informative.

LEP responses from the vertex recording site were pooled across all animals to form a cross-subject prediction. The decision tree classification algorithm was provided with an array of features found to be modulated by behavioural response, including the voltage values at different latencies from laser stimulation (averaged over 30ms bins), the peak amplitude and latency of the LEP, and the change in power in the delta, theta and gamma ranges. The algorithm was trained to distinguish between behavioural scores in three grouped categories corresponding to the ‘no response’, ‘flinch’ or ‘withdrawal’ groupings. To train the model, a training dataset was created using equal numbers of LEPs from each behavioural category. This training set was split into subsets each containing 20% of the data. On each training run, four of these subsets (80% of the data) were used to the train the classifier, with the remaining 20% used to test performance. This procedure was repeated 5 times, using a different 20% as the test set on each run. Accuracy (the proportion of events correctly classified) was evaluated using performance on the test sets (‘5-fold cross validation’). Using this approach, training was performed with between 1 and 10 decision branch splits in the tree, with overall performance reported for the best performing number of splits. Once chosen, this parameter value was then used to train the final model using all the data.

Using features extracted from LEPs with standard pre-processing (bandpass filtering and removal of extreme outlier values, as detailed in Methods) the decision tree classified behavioural outcomes with a 71% accuracy using only two key decision points: peak amplitude and peak latency. To reduce the random variance in the recordings, individual LEPs were then wavelet denoised using the method of Ahmadi et al ^27^ (examples shown in Figure 3a). This technique allows extraction of LEP-related features (described by their wavelet coefficients) from the background noise of the ongoing EEG. This process resulted in a relatively small increase in classifier performance to 73% (using the same process described above for the raw data; the final trained model and performance statistics are shown in Figure 3b and c). However, after wavelet denoising, the optimal classifier decision tree (Figure 3b) was found to use three features – peak amplitude, latency and the value of the voltage at t=210-240ms (likely because the voltage values became more informative after denoising).

**Figure 3.**
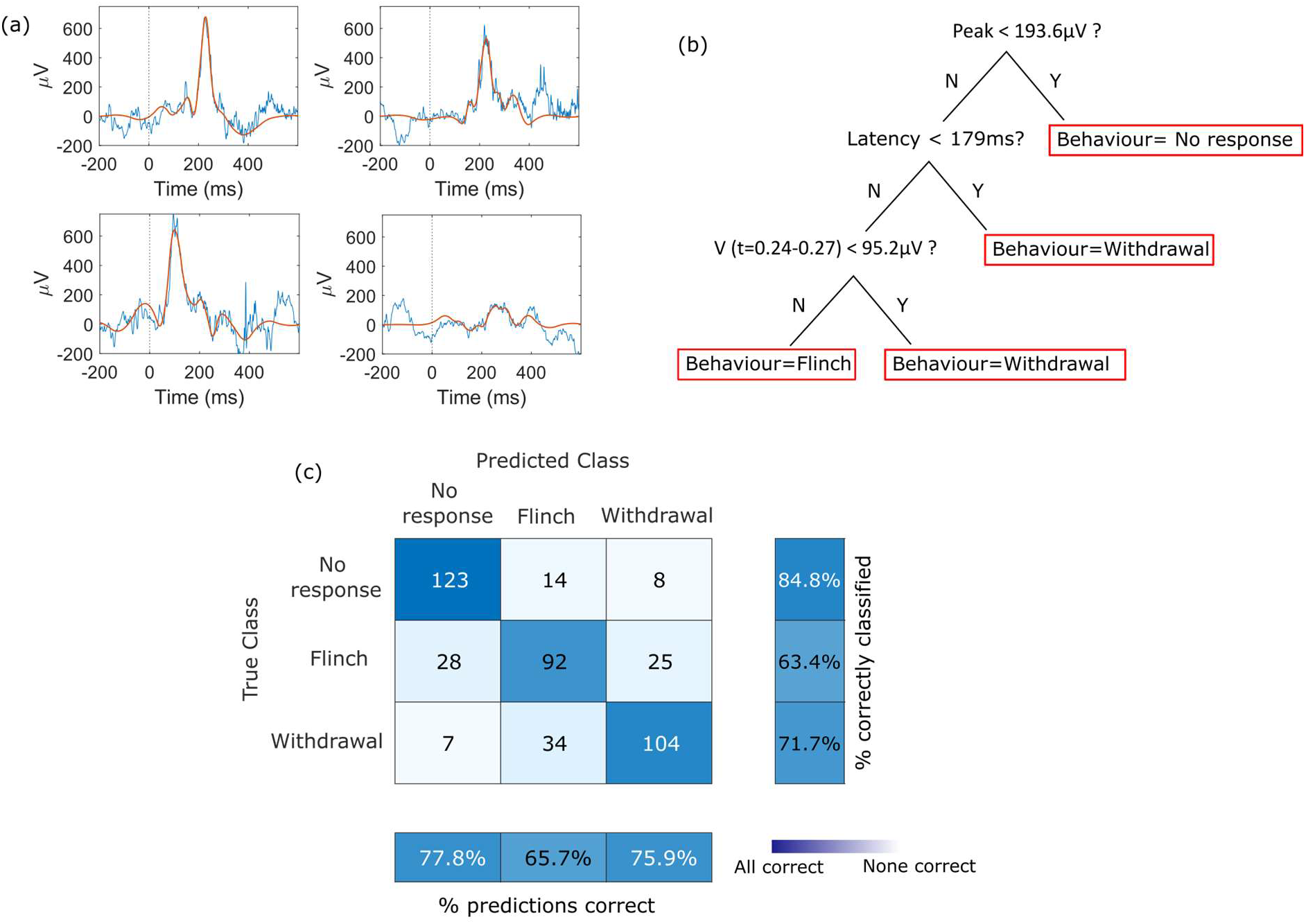
Decision tree classification of individual LEPs. (a) Examples of raw (blue) and denoised (red) single LEPs. (b) Final trained coarse decision tree model for classifying individual LEPs by behavioural response. (c) Matrix illustrating key performance indicators of the coarse decision tree, from 5-fold cross validation training (see main text). Raw values show the number of trials in specific combinations of predicted/true results, e.g. in the top middle square, there were 14 examples of events which were predicted to be in category ‘flinch’ but were actually in category ‘no response’. Summary statistics around the matrix represent the percentage predictions that were correct (bottom) and the percentage of actual results which were correctly classified (right).

To put the performance of the decision tree into context, a selection of other machine learning classifiers were also trained on the denoised data, including fine decision trees, k-means clustering and ensemble techniques. These exhibited comparable performance to the decision tree (61%-75%; see supplementary table 3), however, these approaches provide limited insight into the features used to arrive at the classification and so were not explored further.

The importance of the individual features used by the optimal decision tree classifier were explored in more detail (full results in supplementary table 4). When LEP amplitude was used as the sole feature predicting behaviour accuracy dropped to 63%; despite being permitted up to 5 splits, the classifier was not able to predict any events as belonging to class 2 (‘flinch’), but placed every LEP into either ‘no response’ or ‘withdrawal’. This demonstrates the importance of the latency information in the performance of the classifier. This result was unexpected as the variance of the peak latency with behaviour in the averaged data is insignificant, suggesting that latency would not contribute any extra information when peak amplitude (which does vary strongly with both behaviour and laser energy, Figure 2e and g) is already included.

When laser energy alone was used to predict behaviour (again using a decision tree), the performance of the classifier dropped to 55.4%. When laser energy was included alongside the full set of features used above, both the accuracy of the classifier and the features used were unchanged from previous results (73%, using peak amplitude, latency and voltage at t=210-240ms as features). This confirms that the LEP contributes information which could not be inferred from laser energy alone. Conversely, when the same approach was used to predict laser energy from LEP features, the success rate dropped to 54.0% using decision trees and 46.4-62.1% using other techniques (supplementary table 3) again indicating that additional information is present in the LEP beyond simple noxious stimulus intensity.

Finally, when the classifier was asked only to distinguish between no/low pain-like behaviour (scores 0-2) or pain-like behaviour (scores 3-4), then performance improved to 86.2%. This compares favourably with previous studies, for example, Huang et al ^28^, in which a Naïve Bayesian classifier was used to predict binary pain/no pain states across individuals with a success rate of ~80%.

This machine learning analysis was repeated using features from the EEG recording sites over left sensory cortex and left prefrontal cortex (supplementary table 4; also including data from pairs of sites). The resulting classifiers performed either comparably, or worse than the results from the vertex site alone.

### Bi-modal distribution of latencies in single LEP responses

To better understand the criteria used for decision making by the classifier, the two principal features (vertex peak amplitude and latency) were plotted individually for each LEP event using the denoised data (Figure 4a-c). The peak amplitudes were generally low (<200μV) in the no response behavioural category, thus the classifier is able to categorise low amplitude events as likely belonging to the no response behaviour (first decision point). This representation of the data also reveals two distinct latency groupings for the remaining events, which is particularly apparent within the ‘withdrawal’ behavioural group (Figure 4a). This bimodal distribution is also visible at higher laser energies (Figure 4b). The classifier identifies the short latency/high amplitude events occurring before ~180ms, which are almost uniformly in the withdrawal behaviour category. In contrast, the long latency, high amplitude events are found in both the flinch and withdrawal behaviour group, an area which confuses the machine learning classifier and leads to poorer performance.

**Figure 4.**
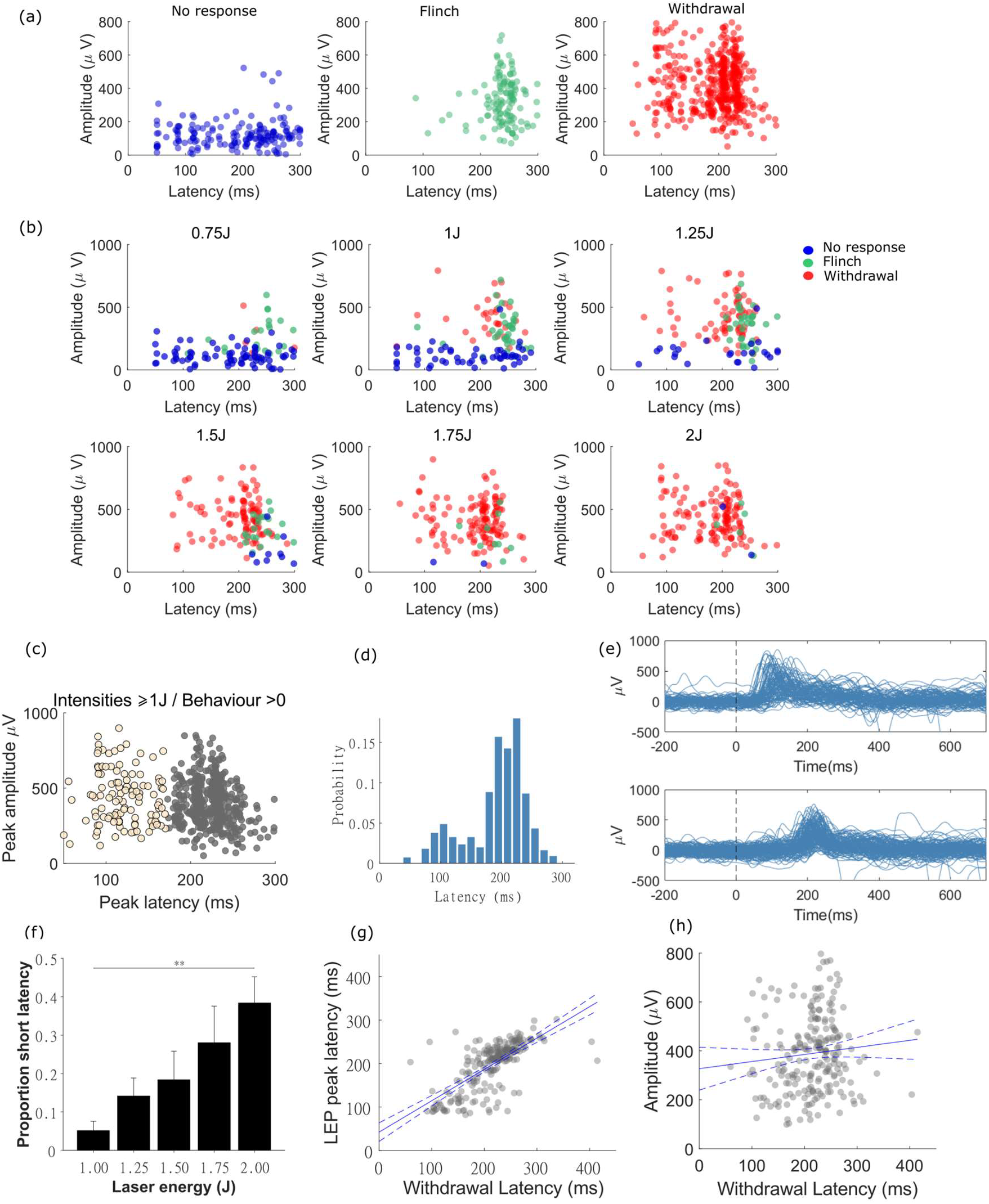
LEPs grouped by latency and amplitude. (a) denoised LEP peak amplitude and latency plotted by behaviour, and (b) laser energy (behavioural response indicated by colours, as used in (a)) showing the appearance of two groups at higher intensities/behavioural responses. (c) LEPs at energies ≥1J and behavioural scores >0, can be split using k-means clustering to form two groups (d) a bimodal distribution is apparent in the probability histogram of latencies for this set reflecting the presence of responses with short and long latencies centred at ~120 and ~220ms (e) 150 randomly selected examples of raw (non-denoised) LEPs identified as short (upper) or long (lower) latency using denoising and k-means clustering. Note that both groups have single peaks at either short or long latency and not both. (f) The proportion of short latency events increases with increasing laser energies (Repeated-measures Friedman test, Dunn’s correction). (g,h) Relationships between withdrawal latencies and LEP latencies/amplitudes, with regression lines and 95% confidence intervals. ** p<0.01.

Further detailed analysis of the two latency groups was conducted on the LEPs where the animal showed a flinch or withdrawal response (using laser energies of 1 J and above, Figure 4c-f). K-means clustering of the LEP data (on the basis of amplitude and latency) identified two groups with short (group 1) and long latency (group 2) peaks. Short and long latency groups had mean peak latencies of 121±3 ms and 223±1 ms respectively and were separated at a latency mid-point of 172 ms (Figure 4c,d). Plotting single trial LEP waveforms from each group further highlights that LEP peaks do not fall onto a continuous spectrum of latencies but consist of two distinct groups (Figure 4e; supplementary figure 3). The proportion of short latency events is greater at higher laser energies (Figure 4f; Friedman test χ^2^(4)=, p<0.01), suggesting that the short latency events are more likely to be evoked by higher skin temperatures.

The frequency of double-peaked LEPs was also investigated (i.e. LEPs containing both short and long latency events), in case peaks at both latencies were present in individual events but had been overlooked by the analysis above. Each LEP was scanned for multiple peaks occurring between 25 and 350ms, subject to the requirement of a minimum distance between individual peaks of 50ms and a minimum peak prominence (i.e. amplitude above baseline) of 100μV. When multiple peaks were detected, the maximum peak was first identified. The LEP was then flagged as a potential double peak if any of the other peaks came within 75% of the maximum peak amplitude. Just 7% of LEP events fell into this category, indicating that the large majority (93%) of LEPs consist of a single peak.

The presence of these latency groupings was not readily apparent within the averaged LEPs (Figure 2b,c). This is partly because the shorter latency events occur less frequently and therefore appear only as a ‘shoulder’ to the left of the main peak of averaged LEPs at the highest laser intensities. Importantly, an exploratory analysis showed that there was robust positive correlation between the hind paw withdrawal latencies and LEP latencies (Pearson’s r=0.7, p<0.001, Figure 4g), suggesting that there is a common mechanism underlying both endpoints (as assessed from the video data e.g. Supplementary Video 1). There was no significant correlation between paw withdrawal latency and LEP amplitude (Figure 4h). This provides further validation of the relevance of LEP latency to the pain-like behaviour.

### LEP latency differences reflect Aδ- vs C-fibre transmission of nociceptive information

We hypothesised that the two latency groups could reflect the mode of transmission of nociceptive information from the periphery. Specifically, the two different latencies may indicate transmission via fast, myelinated Aδ-fibres, or the (more frequent) activation of slower, unmyelinated C-fibres. Previous studies in rats have suggested that whilst C-fibre transmission of LEPs is more frequently observed, Aδ-mediated events are also seen^17,19,20^. If this is the case, the differences in peak latencies between LEPs mediated by Aδ or C-fibres should reflect the different speeds of transmission along these two fibre types: ~10ms^-1^ for Aδ-fibres, ~0.75ms^-1^ for C-fibres^29^. By assuming a 10cm distance along the leg of an adult rat then the transmission latency will differ by ~120ms – similar to the ~100ms difference between the mean latencies of the two groups of LEPs (Figure 4c).

In order to explore this hypothesis further, we sought to inhibit the C-fibre events using capsaicin, a TRPV1 agonist which ‘defunctionalises’ these axon terminals at high concentrations^30,31^, with the prediction that putative short latency, Aδ-fibre mediated LEPs would be left intact (specifically, via capsaicin insensitive Aδ mechano-heat fibres type 1 [AMH-I]^17,32,33^). LEPs were recorded before and 4, 28 and 52 hours after intra-plantar capsaicin injected subcutaneously to the hind paw. A reduction in pain-like behaviour was seen following stimulation of the capsaicin-treated paw (n=6 responder animals, Figure 5a,b; supplemental figure 4b). Because the behavioural responses were stable across all post-capsaicin timepoints (Supplementary Figure 4a), their data were pooled into a single post-capsaicin dataset. The reduction in pain-like behaviour was also reflected in a reduction in the amplitude of the average LEP from all stimuli (supplemental figure 4c), but with an apparent preferential loss of the peak and retention of the early shoulder. To analyse this effect further we studied the post-capsaicin events that generated a behavioural flinch or withdrawal response. The features of these LEPs were markedly different relative to the pre-capsaicin set (Figure 5c), with a distinct peak appearing at the shorter latency of ~100ms and a corresponding proportional increase in short latency (t<172ms) LEP peaks from a mean of 19±6% to 43±13% on the treated side (Figure 5d). This was reflected in a commensurate reduction in overall mean ERP latency (from 206.6±6.3ms before to 161.1±9.5ms after capsaicin, p<0.001, Figure 5e). This suggests that the predominant mechanism underlying LEP generation was changed post-capsaicin administration; it is likely that a greater proportion of the remaining withdrawal responses were associated with Aδ-fibre mediated LEPs.

**Figure 5.**
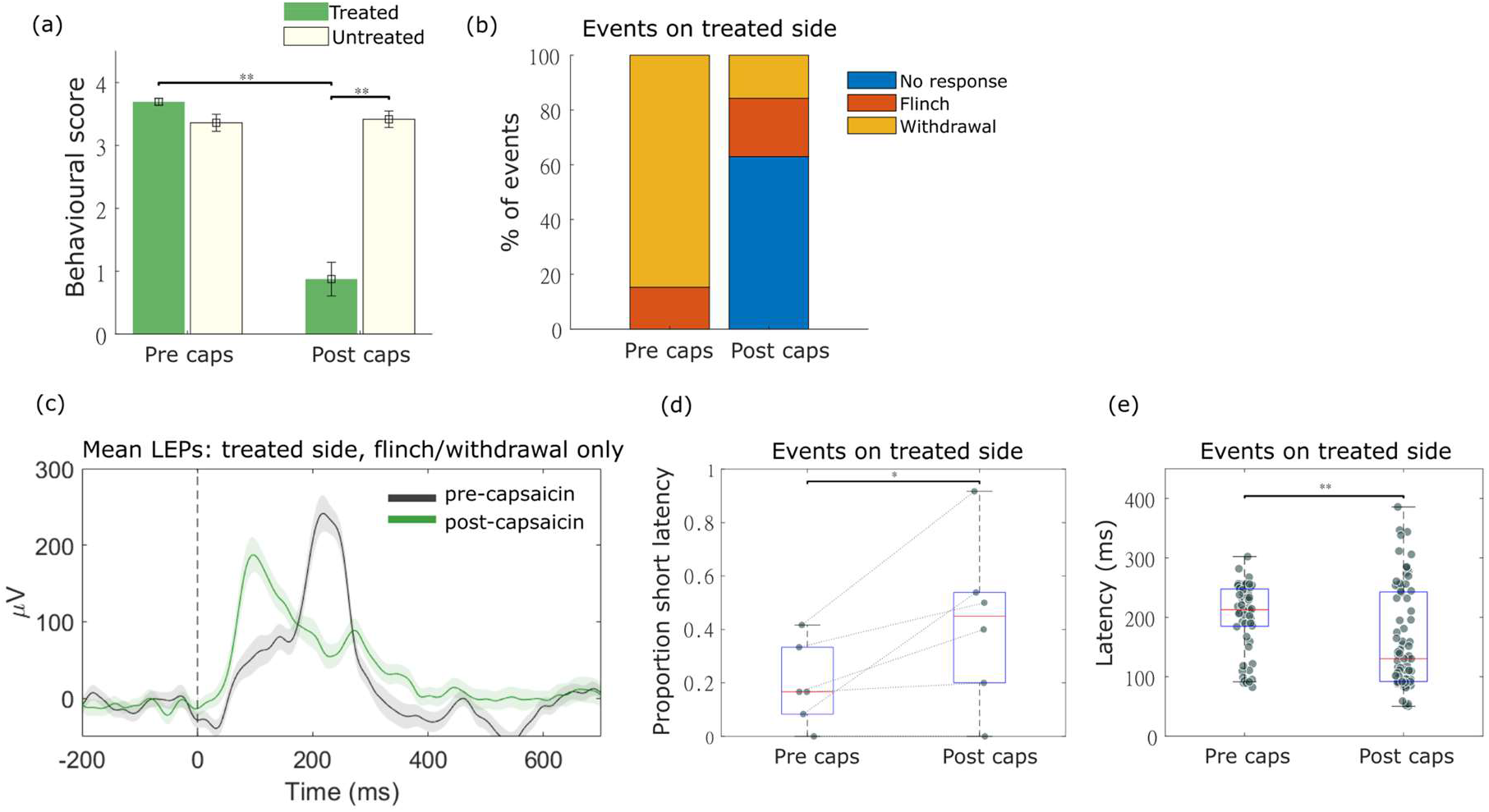
LEPs and capsaicin. (a) behavioural scores following 1.5J laser stimulation of the capsaicin-treated foot (green bars) and the untreated foot (blue bars), repeated measures ANOVA with time and side as within subject factors, Bonferroni correction for repeated comparisons. (b) Capsaicin caused many more ‘no response’ events. (c) The mean raw (not denoised) LEP before and after capsaicin restricted to trials where a behavioural response remained, shows a flattening of the original peak at ~250ms and introduction of a prominent peak at ~100ms post capsaicin. (d) Mean latency of LEP peaks for each animal, before and after capsaicin. The proportion of short latency (peak latency < 0.172s) events showed a significant increase post-capsaicin (Lilliefors test for normality, paired t-test. (e) LEP peak latencies for all events, not grouped by animals, showing a significant reduction in overall latency, and a greater proportion of events at shorter latencies (as visible in (c), Lilliefors test for normality, Wilcoxon rank-sum test).

## Discussion

Our starting point for this study was to ask whether the rat LEP had validity as a translational biomarker of pain and to identify the features that could be most informative. The results from our study suggest that variations in LEP features, such as peak amplitude, reflect pain-related behaviour in naïve rats. In particular, the amplitude of the main vertex peak at ~250ms in averaged LEPs correlated strongly with behavioural response – even when laser energy was held constant. This indicates that the LEP encodes aspects of the decision-making process around pain-like behaviour rather than being a simple proxy of stimulus intensity or sensory input. This is consistent with the conclusions of previous studies in both humans and rodents^12,34^.

We deployed a machine learning approach to ask whether this relationship could be used to predict behavioural response from single-trial EEG data. A coarse decision tree classifier was able to predict the behavioural response associated with individual LEPs to an accuracy of 73% (with particular success in discriminating no response from withdrawals (~90% accuracy)). We found the optimal classifier used both amplitude and latency features of single LEPs. It was apparent that the latency of LEP peaks followed a bimodal distribution - with around ~30% of events occurring at a shorter latency of ~120ms, a result not initially evident in the averaged LEPs. This indicates that when averaging takes place, meaningful differences in the form of LEP responses to individual stimuli are lost. This is particularly the case when only a subset of events – here, the short latency events – differ in morphology. The appearance of this set of LEPs at the higher laser intensities/behavioural responses manifests only as a small ‘shoulder’ on the side of the main peak in averaged data (visible in Figure 1d and e). Interestingly the inclusion of LEP data from other EEG recording sites or frequency power spectrum data did not improve the performance of the classifier, indicating that they did not carry additional information that was better able to discriminate between behavioural responses. The main area where the classifier performance could be improved is in distinguishing between flinch and withdrawal responses, and future studies will be needed to assess whether this could be better achieved for example by the inclusion of local field potential data from sites believed to encode pain intensity and aversiveness such as the insula or amygdala whose activity is not captured in surface EEG recordings.

As predicted based on LEP studies in humans^35,36^, peripheral application of capsaicin (causing defunctionalisation of C-fibres^30^) resulted in a large reduction in the amplitude of the LEP. This was accompanied by a significant reduction in the number of withdrawal responses to a given intensity of laser stimulation. These effects were predominantly driven by a reduction in putative C-fibre responses, with Aδ responses largely spared (and therefore contributing a greater proportion of the remaining responses). It is likely that capsaicin-insensitive AMH-I fibres are a significant contributor to these remaining events, however the majority of the remaining responses are still in the C-fibre latency range. Comparable studies in humans have typically observed complete loss of LEP responses, however these studies used multiple rounds of capsaicin application over several days to ensure near-complete denervation of the epidermis^35,36^, a difference which may explain the small number of responses remaining here (where a single capsaicin application was used). Studies in humans have also demonstrated that capsaicin-induced desensitisation of Aδ-mediated, laser-evoked responses is restricted to the area of capsaicin application, whereas C-fibre desensitisation covers a larger area, likely due to the larger receptive fields of C-fibres^37^. Consequently, some of the remaining Aδ-mediated responses in rats may also be due to inadvertent stimulation of fibres that were not directly exposed to capsaicin, a phenomenon that is much less likely for C-fibres. The finding that both Aδ and C-fibres are likely to contribute to the LEP in rodents is in contrast to previous studies using averaged data, which have concluded that in rats LEPs are mediated only by C-fibres^11,13^. However it is in agreement with a number of studies using a range of methodologies that have indicated a role for Aδ-fibres in thermal nociception in rodents^17–22^. Our results suggest that the latencies of individual LEP peaks convey information about the mode of transmission of nociceptive information from the periphery.

Interestingly there appeared to be a mutual exclusivity in the single LEP responses with either a short latency or a long latency peak (an effect seen both within and across animals) with little evidence for peaks at both Aδ and C-fibre latencies (likely overestimated at ~7% of all LEPs due to inclusion of some peaks that were the product of noise). The size of the illuminating laser spot is relatively large (4mm diameter) meaning it is unlikely to be a simple case of stochastic activation of one afferent terminal class or the other based on small variations in the location of stimulation. Rather it is likely consistent with the observation that the threshold for thermal activation of Aδ-fibres is higher than that of C-fibres^19–21^, and so are less likely to be activated. However, when Aδ-fibres are engaged their activity precedes and is powerful enough to dominate the C-fibre nociceptive barrage and drive behaviour and the LEP. A similar bimodal distribution of short and long latency withdrawals (presumed Aδ and C-fibre mediated) has been noted in mice with selective optoactivation of classes of primary afferent, and particularly relevant to our study, to those expressing TRPV1^38^. Browne & colleagues^38^ suggested that the two subsets of behavioural response were due to intrinsic properties of the afferents and their transmission pathways in the CNS, rather than reflecting differences in the transduction mechanism. It has also been proposed that Aδ input may transiently inhibit C-fibre mediated nociceptive drive on to spinothalamic tract neurons^39^. This may account for the observation in humans that a C-fibre LEP is only seen when the Aδ LEP is blocked^1,15^. Alternatively, similar occlusion of the human ultra-late C-fibre mediated potential by Aδ activation has been suggested to be due to a cortical refractory state ^1,2,40^. While the precise mechanism is uncertain based on our studies, this property again emphasises the cross species similarity in processing. We speculate this mutual exclusivity in the circuit organisation could act to prevent the generation of two sequential motor withdrawal responses to a given stimulus which would be unlikely to convey an advantage and may impair/delay co-ordinated locomotor escape behaviour.

Our findings extend the potential translational validity and utility of rodent LEPs by demonstrating the presence of both C- and Aδ-fibre mediated responses in conscious behaving animals. This has previously required the use of anaesthetised preparations where a carefully graded heat stimulus could be delivered. Indeed, we note that the ability to discriminate between these two pathways of nociceptive transmission by using single-trial LEP analysis increases the level of mechanistic insight. This may allow the profiling of pharmacological activity in a fibre-specific manner, adding additional evidence of target engagement, as was demonstrated here for topical capsaicin. As another example, analgesics targeting the Nav1.8 channel^41,42^ (found predominantly in C-fibres^43–45^) would also be predicted to reduce the C-fibre component of remaining LEPs, whilst leaving the Aδ component intact.

In summary, the findings described here indicate that rat LEPs have a set of characteristic properties that support them being a useful, translatable measure of pain. Furthermore, adoption of the single trial analysis, machine learning and wavelet filtering approaches may help to identify further novel and important mechanistic features of interest across species.

## Materials and Methods

### Animals

All experiments were performed in accordance with the United Kingdom Animals (Scientific Procedures) Act 1986 with approval from the United Kingdom Home Office and local Animal Welfare and Ethical Review Board (Eli Lilly). Studies were conducted on a total of 12 adult, male Wistar rats (345-387g on date of surgery) who were individually housed on a 12 hour light/dark cycle with food and water provided ad libitum.

### Surgical implantation for EEG recordings

Each rat was anaesthetised with isoflurane and implanted with an array of four stainless steel EEG skull screws (00-96×1/16 Plastics One, USA). The four recording positions were located over: motor cortex (AP +3.5mm, ML −2.0mm); vertex/cingulate cortex (AP −1.5mm, ML 0.0mm); primary somatosensory cortex - hindlimb region (AP −1.9mm, ML −2.6mm) and visual cortex (AP −6.6mm, ML +4.2mm) [all measurements relative to Bregma]. In addition three depth electrodes (200μm insulated silver wire, Advent Research Materials Ltd, UK) were inserted to: insular cortex (AP +2.8mm, ML +3.9mm, DV −3.9mm); infralimbic cortex (AP +2.7mm, ML +0.6mm, DV −4.2mm) and Amygdala (AP - 1.9mm, ML +4.0mm DV −7.5mm) [AP/ML measurements relative to Bregma, negative ML values represent Left side, DV measurements relative to brain surface]) connected to a 12 channel circular connector (Omnetics Connector Corporation, USA). An additional screw was placed in the occipital bone and acted as the ground electrode. Note that the EEG site over motor cortex and the depth electrodes were not used in the analysis above. Light-curable composite (Revolution Formula 2, Kerr Dental, USA) was used to bind the implant to the skull. The analgesic Carprofen (5mg/Kg SC) was administered at the end of surgery and on the morning of the first postoperative day, with subsequent analgesia provided by Meloxicam (1mg/kg PO) on each of the first 7 days post-surgery. The prophylactic antibiotic (Sulfatrim PO [4.8mg/Kg Trimethoprim & 24mg/Kg Sulfamethoxazole]) was administered twice daily from the morning of surgery to 7 days postoperatively. Rats were given a minimum period of 14 days following surgery before undergoing further experimental procedures.

### Experimental protocol

The experiments were performed in a custom-built enclosure (Apogee Engineering Analysis Solutions Ltd, UK), consisting of an upper chamber with four individual compartments for rats, and a lower chamber where the fibre-optic cable from an 1340nm wavelength Nd:YAP laser (Stimul 1340, Electronical Engineering Group, Italy) terminated. The floor of the rodent compartments was constructed of borosilicate glass, through which the laser beam could pass. The primary Nd:YAP laser could be targeted by visualising a low power laser (635nm wavelength) on the under-surface of the rat using a USB camera connected to a PC. The position of the laser was adjusted using a joystick-controlled motorised xy-stage (Igus GmbH). For all experiments, the laser stimuli (diameter: 4mm; duration: 4ms) were delivered to the centre of the plantar surface, alternating between left/right hind-paws (interstimulus interval: >30s).

Rats were habituated to the apparatus for 5 days before the stimulus response protocol commenced. The protocol was split across 4 recording sessions, with an inter-session interval of at least one day. Within each session, rats were exposed to 18 laser stimuli at 6 different intensities (0.75, 1.00, 1.25, 1.50, 1.75 & 2.00J). Stimuli were presented in blocks of 3 of the same intensity according to a balanced design. At the end of each session, a further 3x 2J stimuli were applied to either the dividing walls between the rats (sessions 1 & 2), or to the ceiling of the enclosure (sessions 3 & 4) as a control for potential auditory evoked responses. EEG data was acquired at a sampling rate of 19.525kHz (filtered 0.35 - 9700Hz) using wireless TAINI transmitters (TainiTec Ltd, UK)^48^.

For the exploratory capsaicin experiment, 10 animals retained high quality EEG data and were included in the study. Recordings were performed across four days, with two sessions per day, four hours apart. During each session, rats were stimulated by the laser on the plantar surface of the hind paw (12 stimulations per paw, alternating left/right) at an energy level of 1 and 1.5J in sessions 1 and 2 respectively. Immediately prior to the first recording session on day 2, rats were injected subcutaneously with capsaicin (20μg in 100μl DMSO) in the plantar surface of the hind paw (randomly allocated to left/right paw in a balanced manner). Five of the rats showed evidence of defunctionalisation with a reduction in behavioural responses to laser stimulation (1.5J) and were included in this exploratory analysis.

### Behavioural scores

Behavioural responses were recorded for each stimulus application by the experimenter based on the following scale: 0: no response, 1: flinch (or some sign of awareness of the stimulus), 2: transient foot lift, 3: withdrawal and large body movement, 4: withdrawal and licking.

A repeated measures ANOVA was used to analyse the relationship between mean behavioural score and laser energy (with energy and side of stimulation as within-subject factors and mean behavioural score calculated for each animal at a given laser energy/side).

### Pre-processing of EEG

EEG was processed and analysed using custom MATLAB scripts. Missing samples in the EEG data caused by errors during wireless transmission were linearly interpolated in MATLAB and the resulting signal high-pass filtered (zero-phase offset, 2nd order Butterworth filter; cut-off 0.35Hz) to remove the DC offset. Signals were then low-pass filtered (zero-phase offset, 2nd order Butterworth filter; cut-off 250Hz) and segments of data ±5s around each laser stimulation extracted. Each window of data underwent further basic pre-processing to remove noise and artefacts. Samples over a threshold level of 750μV were removed and replaced with linearly interpolated values using MATLAB’s built in interpolation function ‘interp1’. Trials where there was more than 0.1s of interpolation, or where there were more than 10 discrete segments of interpolation in the period immediately adjacent to the laser stimulus (−0.5 to 0.7s) were excluded. Each event was baseline normalised by subtracting the mean EEG voltage from the 0.5s preceding the laser stimulation from all samples.

For analysis of the LEP peak features, data was additionally downsampled by a factor of 5 (from 19525Hz and low pass filtered with a cut off of 30Hz using a zero-phase offset, 3^rd^ order Butterworth filter (built in MATLAB function ‘butter’). For analysis of power in the delta (0.5-5Hz), gamma (50-100Hz) and theta (5-12Hz) frequency ranges, raw data was bandpass filtered using a 2^nd^ order Butterworth filter; the power was calculated using a Hilbert transform. Baseline power was calculated as the mean of the power in the 2 seconds prior to the laser event.

### Analysis of average LEPs

Mean LEPs were calculated using all events in a specified category (i.e. with a given laser energy or behavioural response). This averaged LEP was then used to extract the peak amplitude and latency. The peak was identified as the maximum point occurring within an event window of 0.05-0.35s following the laser stimulus. One way repeated measures ANOVA were used to analyse the variation in peak amplitude or latency, using either behavioural score or laser energy as within subject factors. For comparisons using either fixed behavioural response or laser energy; the laser energy was grouped for analysis into three categories to match the three categories used for behaviour; these were 0.75-1J, 1.25-1.5J, 1.75-2J.

For calculations of power spectrum characteristics, the change in power over the event window (relative to the mean over a 5 second baseline) was calculated for each individual laser stimulus. Power curves were then averaged within categories (i.e. for a given animal and laser energy / behaviour). The peak and latency of these curves were then analysed in the same way as the LEP peaks.

### Potential for laser generation of auditory evoked potentials

It has been reported that rodent LEPs can be contaminated by fast auditory evoked potentials (AEPs)^11^. These are proposed to result from ultrasonic sounds generated by the rapid skin heating caused by laser stimulation). To test for the potential contribution of AEPs to the LEP waveforms, EEG responses were also recorded while the laser was targeted at the base of the dividing walls separating the rats. This generated obvious AEPs which were clearly distinct from the paw stimulus triggered LEPs in both timing and morphology (supplementary figure 1, the grand averaged AEP has a small positive peak at ~50ms, of amplitude 100μV – this clearly precedes the LEP peaks found at 200ms, Figure 2a-d). This provides confidence that recorded LEPs were not contaminated by (or confused with) AEPs.

### Analysis of single trials and use of machine learning algorithms

Single trial LEPs were denoised using the EP_DEN software described in Ahmadi et al ^27^. Briefly, the method uses a dataset of multiple event related potentials (ERPs - here these are LEPs) to calculate the coefficients of wavelet components relevant to the post-event ERP, using a baseline period as a comparator. Individual ERPs provided to the software are then reconstructed using only these ‘informative’ wavelet components, removing background noise.

Raw LEPs were resampled from a sampling rate of 19525Hz to 10923Hz using the MATLAB function ‘resample’. This was done in order to make the length of the entire signal equal to a power of 2 (here, 2^15^ samples) as required by the EP_DEN software. To calculate wavelet coefficients, the entire dataset of resampled LEPs across all animals, intensities and behaviours was provided to the EP_DEN software, which returned denoised LEPs, alongside values of the primary peak amplitude and latency for each event. These values of amplitude and latency were then used as part of the feature set to train the classifiers. For single trial binned voltage values, a moving average filter was applied the denoised data using a window of 30ms. Samples of this averaged data were acquired from 30ms bins between 0.03 and 0.55s relative to the laser stimulus. Changes in spectral characteristics for individual events were calculated by bandpass filtering and Hilbert transforming raw data (in MATLAB), then calculation of the amplitude of the change in power relative to baseline. Feature sets were normalised by subtracting the mean and dividing by the standard deviation before being used in training.

Coarse decision trees were trained on the feature set using the built-in MATLAB function ‘fitctree’, with 5-fold cross validation, and a minimum leaf size of 15. The accuracy, confusion matrix (including the recall and precision statistics shown) were extracted from the fitted model and were calculated using only examples which were not used in training. The final model (shown in Figure 3) was trained using all available data. All other machine learning algorithms were trained using built in models in the MATLAB classifier toolbox GUI. When classifiers were trained to predict laser energy, the dataset was combined into 3 groups of intensities (0.75-1J, 1.25-1.5J, 1.75-2J) to create a comparable condition to that used when classifying the 3 behavioural responses.

K-means clustering by peak amplitudes/latencies used the MATLAB function ‘kmeans’, using two groups. The threshold between the two groups was calculated as the midpoint between the longest peak of the short latency events and the earliest peak of the long latency event group.

When calculating the proportion of fast events at each laser energy, only two rats exhibited at least 6 valid responses at 0.75J (e.g. due to noise, or lack of behavioural response), and so this energy level was removed from this analysis due to the inaccuracy of subsequently calculated parameters. Across the remaining energy levels, n=5 (of 12) rats were excluded as they did not exhibit a minimum of 6 valid events at each energy level. Remaining data was analysed using a Friedman test (repeated measures; GraphPad Prism v9.2.0)

## Acknowledgements

The authors wish to thank Professor Luis Garcia-Larrea for providing helpful feedback on the manuscript.

## Competing interests

This project has received funding from the Innovative Medicines Initiative 2 Joint Undertaking under grant agreement No [777500]. This Joint Undertaking receives support from the European Union’s Horizon 2020 research and innovation programme and EFPIA.

The statements and opinions presented here reflect the author’s view and neither IMI nor the European Union, EFPIA, or any Associated Partners are responsible for any use that may be made of the information contained therein.

## Supplementary figures

**Supplementary Figure 1.**
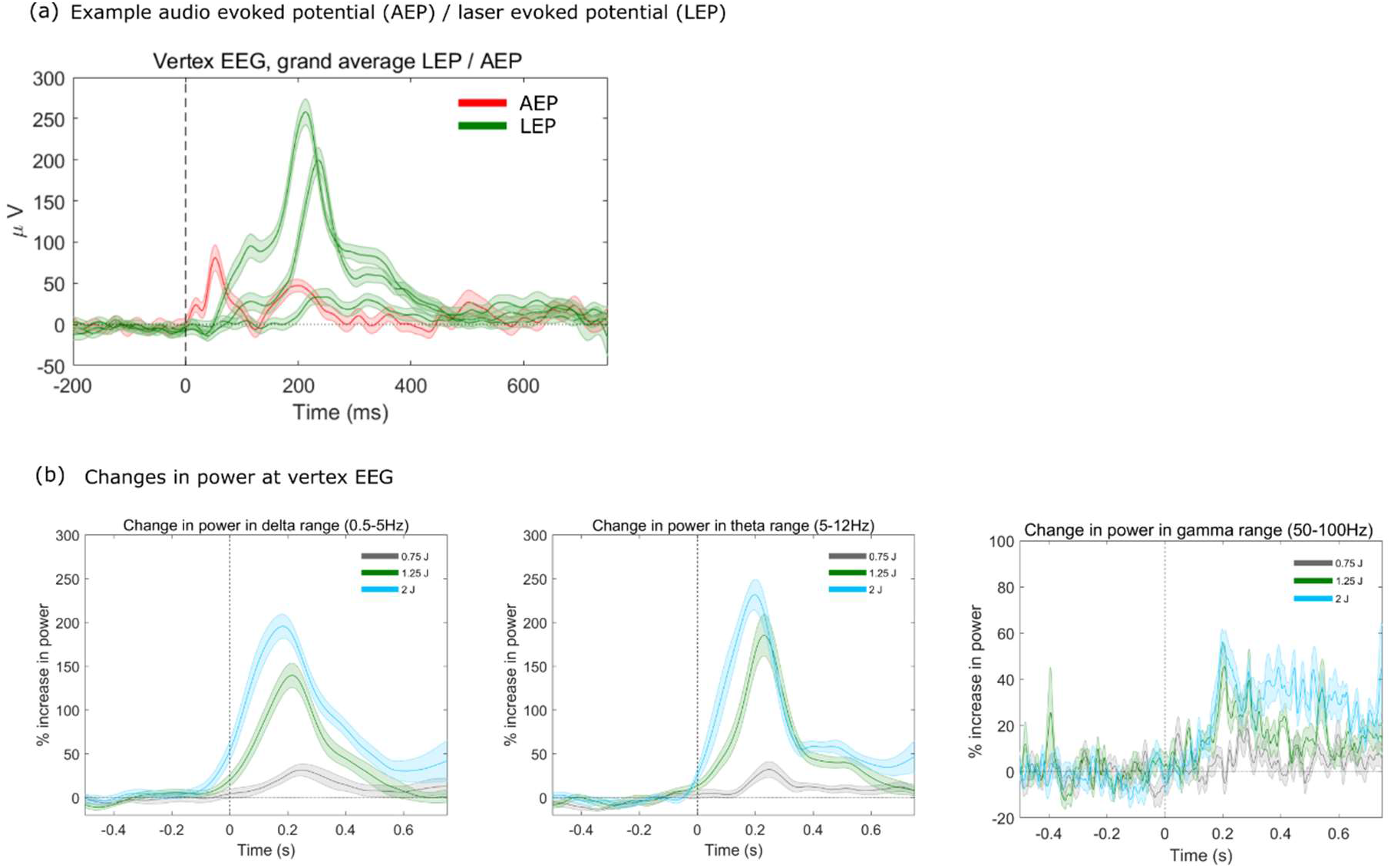
(a) mean auditory evoked potential across all animals, generated by targeting the laser onto a metallic surface near the animal. The form of the auditory evoked potential is distinct from the LEP, with a peak at ~50ms (b) changes in power in the delta, theta and gamma range at varying laser intensities.

**Supplementary Figure 2.**
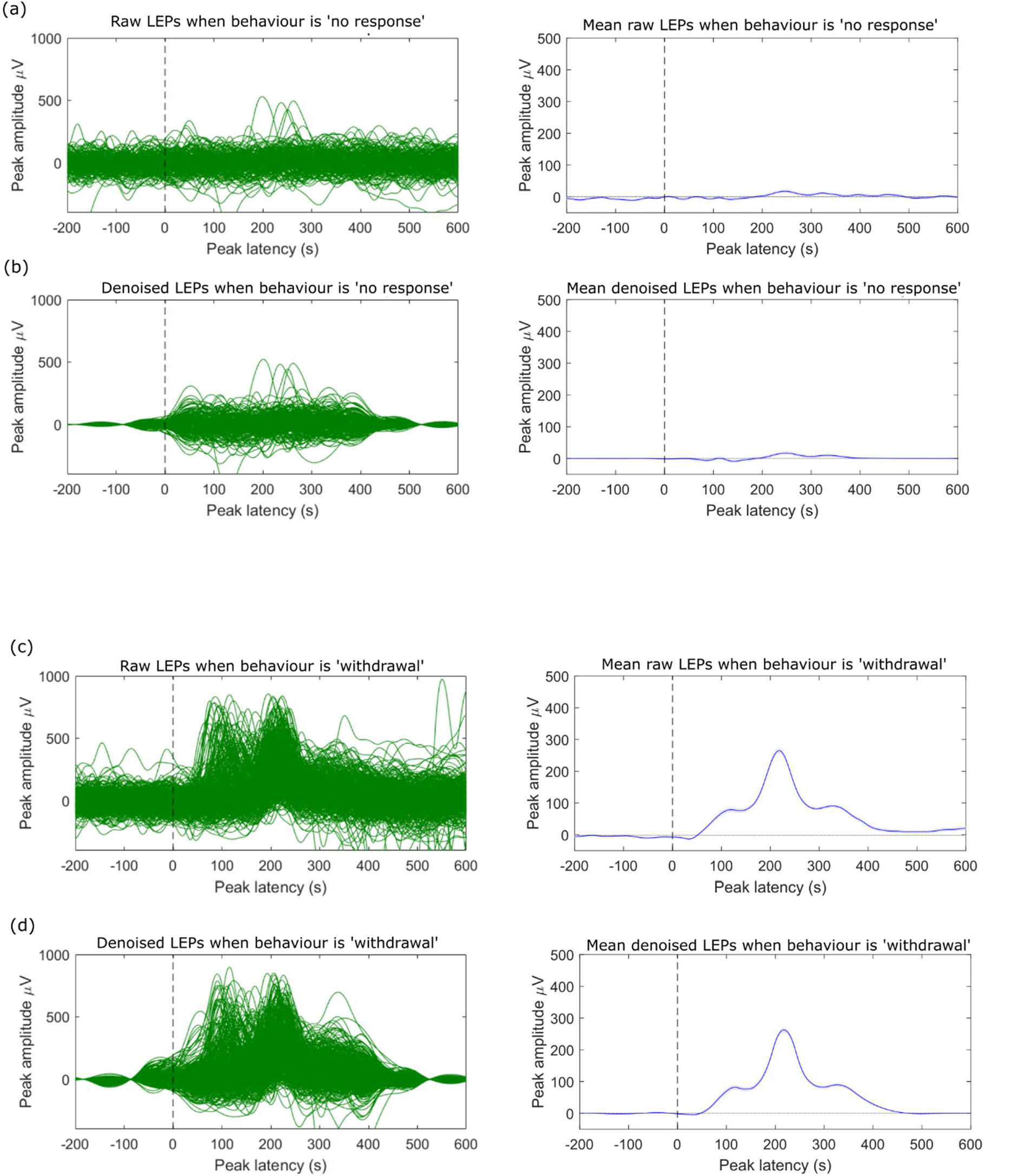
(a-b) For behaviours classified as no response, most individual vertex LEPs do not rise above the noise (left hand plots). Little/no clear peak is apparent in the mean of either the raw or denoised data.(c-d) For the withdrawal responses, peaks rise clearly above the noise in both raw and denoised versions. Note that in the averaged data for these LEPs, the two groups of peaks have merged into one peak with ‘shoulders’ either side.

**Supplementary Figure 3.**
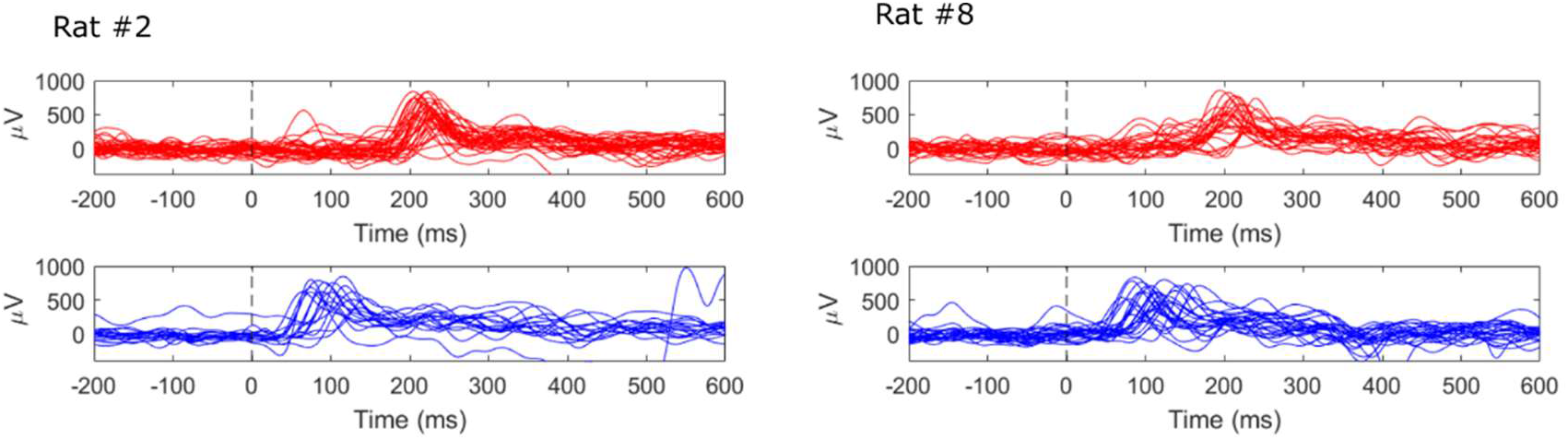
Distinct sets of LEPs at short and long latency are seen within the datasets for individual animals.

**Supplementary Figure 4.**
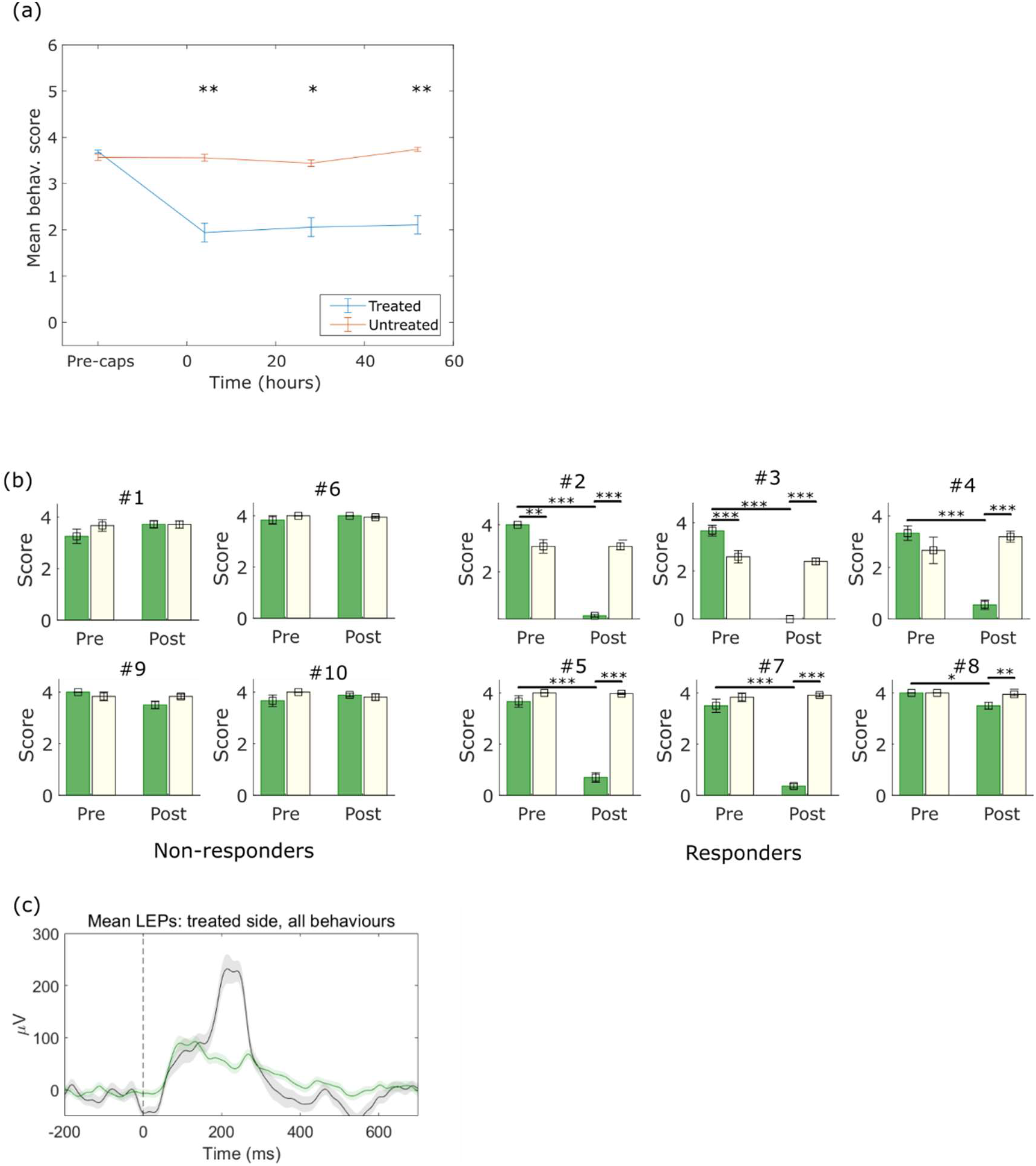
(a) Mean behavioural responses at all timepoints (all animals), pre- and post-capsaicin. (b) Breakdown of behavioural responses by animal, categorized as responders and non-responders. (c) Mean LEP pre and post capsaicin over all behavioural responses for the responder group.

**Supplementary table 1.** Results for additional EEG recording sites. (a) Significance of relationships between LEP amplitude / latencies versus laser energy / behavioural outcome. (b) Significance of relationships between LEP amplitude versus laser energy at a given behaviour / behaviour at specific laser energy (*** p<0.001, ** p<0.01, * p<0.05).

| Site | LEP amplitude versus... |  | LEP latencies versus... |  |
| --- | --- | --- | --- | --- |
|  | Laser energy<br>(ANOVA F, p) | Behaviour<br>(ANOVA F, p) | Laser energy<br>(ANOVA F, p) | Behaviour<br>(ANOVA F, p) |
| Left frontal | 25.8, *** | 46.2, *** | 4.9, *** | 1.5, NS |
| Left sensory | 25.6, *** | 46.7, *** | 2.0, NS | 1.5, NS |
| Vertex | 34.8, *** | 72.3, *** | 2.6, * | 1.3, NS |

**Supplementary table 1. Results for additional EEG recording sites.**
| Site | LEP amplitude versus... |  |
| --- | --- | --- |
|  | Laser energy at fixed behaviour (1-2)<br>(ANOVA F, p) | Behaviour at fixed laser energy<br>(1.25-1.5J) |
| Left frontal | 1.1, NS | 15.8, *** |
| Left sensory | 0.4, NS | 16.4, *** |
| Vertex | 0.8, NS | 31.8, *** |

**Supplementary table 2.**
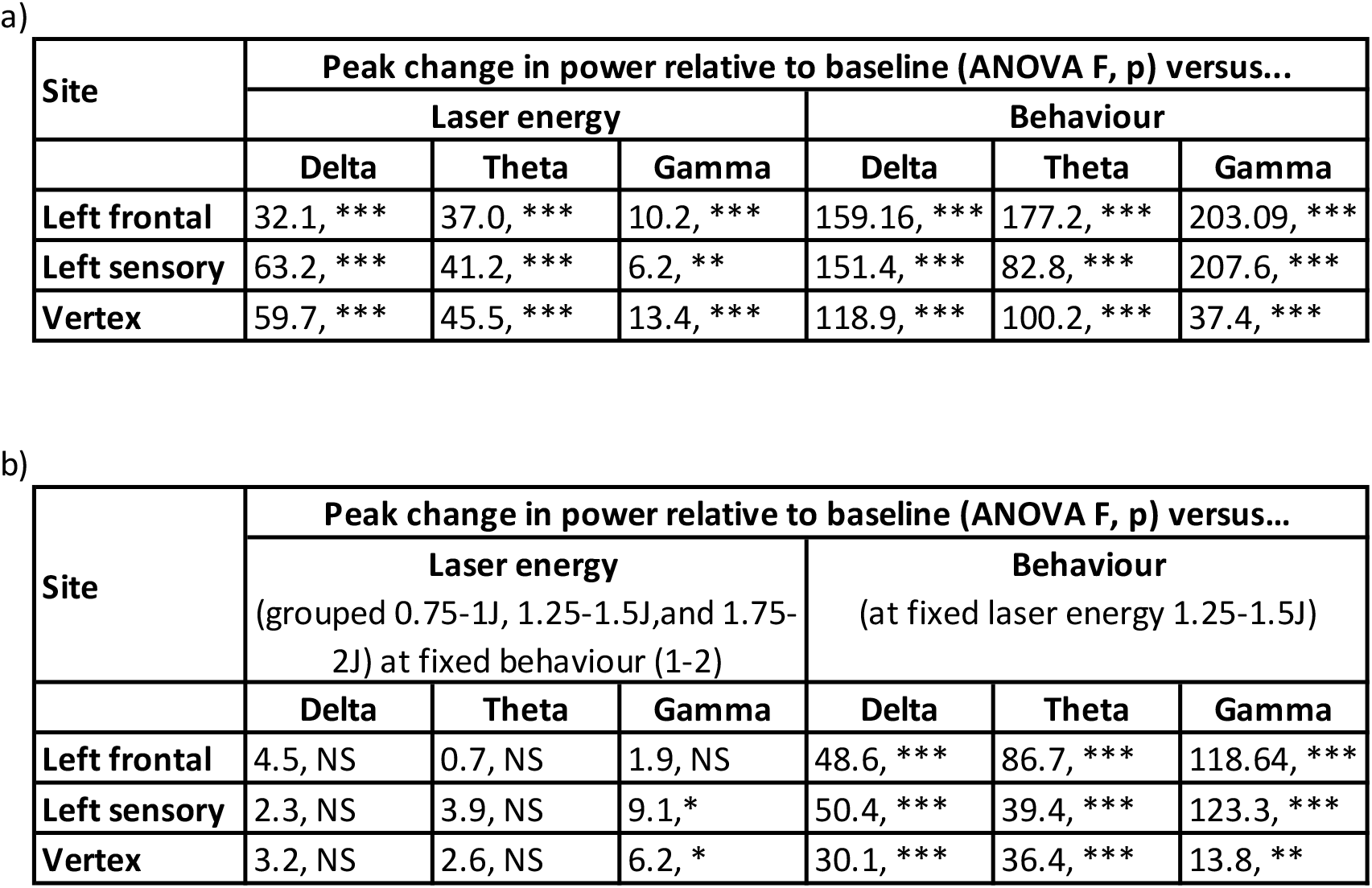
Change in spectral power by site. (a) Significance of relationships between maximum change in spectral power and laser energy / behavioural outcome. (b) Significance of relationships between spectral power and (i) laser energy at constant behaviour, (ii) behaviour at constant laser energy (*** p<0.001, ** p<0.01, * p<0.05).

**Supplementary table 3.** Results from alternative classification algorithms.

| Type | Subtype | Accuracy (% correct) :<br>behaviour classification | Accuracy (% correct) :<br>laser energy |
| --- | --- | --- | --- |
| Decision trees | Coarse | As above (73.3) | 57.7 |
|  | Medium | 70.1 | 56.1 |
|  | Fine | 71.7 | 52.4 |
| Linear Discriminant | N/A | 72.4 | 59.3 |
| Naïve Bayes (Gaussian) | N/A | 68.3 | 57.9 |
| Support Vector Mechanism | Linear | 74.7 | 62.1 |
|  | Cubic | 68.1 | 52 |
|  | Quadratic | 66.9 | 54.5 |
|  | Medium Gaussian | 70.1 | 57.5 |
|  | Fine Gaussian | 52.6 | 46.4 |
| K-nearest-neighbour | Fine | 61.1 | 51.3 |
|  | Medium | 66.7 | 58.2 |
|  | Coarse | 72.2 | 59.1 |
| Ensembles | Bagged Trees | 72.2 | 55.9 |
|  | KNN | 65.3 | 55.6 |

**Supplementary table 4.**
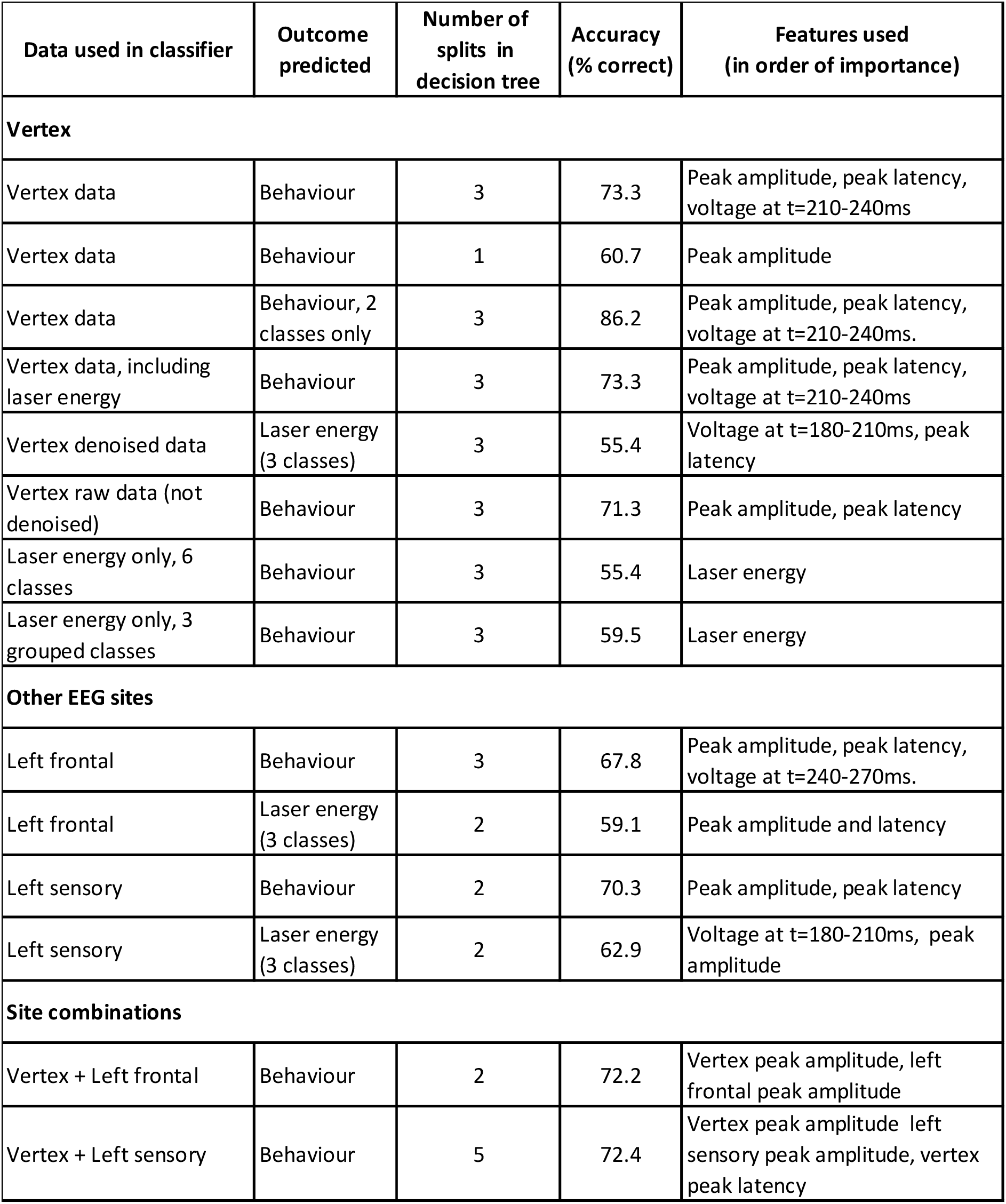
Results from coarse decision tree using alternative features, sites and parameters. Unless otherwise stated: 1) predictive features are the same as used in Figure 3 (i.e. voltage values at varying latency from laser, peak amplitude and latency, changes in power in delta, theta and gamma ranges). 2) Data used is denoised as per explanation in main text. 3) Behaviour was classified into 3 categories (no response, flinch and withdrawal).

| Data used in classifier | Outcome predicted | Number of splits in decision tree | Accuracy (% correct) | Features used (in order of importance) |
| --- | --- | --- | --- | --- |
| **Vertex** |  |  |  |  |
| Vertex data | Behaviour | 3 | 73.3 | Peak amplitude, peak latency, voltage at t=210-240ms |
| Vertex data | Behaviour | 1 | 60.7 | Peak amplitude |
| Vertex data | Behaviour, 2 classes only | 3 | 86.2 | Peak amplitude, peak latency, voltage at t=210-240ms. |
| Vertex data, including laser energy | Behaviour | 3 | 73.3 | Peak amplitude, peak latency, voltage at t=210-240ms |
| Vertex denoised data | Laser energy (3 classes) | 3 | 55.4 | Voltage at t=180-210ms, peak latency |
| Vertex raw data (not denoised) | Behaviour | 3 | 71.3 | Peak amplitude, peak latency |
| Laser energy only, 6 classes | Behaviour | 3 | 55.4 | Laser energy |
| Laser energy only, 3 grouped classes | Behaviour | 3 | 59.5 | Laser energy |
| **Other EEG sites** |  |  |  |  |
| Left frontal | Behaviour | 3 | 67.8 | Peak amplitude, peak latency, voltage at t=240-270ms. |
| Left frontal | Laser energy (3 classes) | 2 | 59.1 | Peak amplitude and latency |
| Left sensory | Behaviour | 2 | 70.3 | Peak amplitude, peak latency |
| Left sensory | Laser energy (3 classes) | 2 | 62.9 | Voltage at t=180-210ms, peak amplitude |
| **Site combinations** |  |  |  |  |
| Vertex + Left frontal | Behaviour | 2 | 72.2 | Vertex peak amplitude, left frontal peak amplitude |
| Vertex + Left sensory | Behaviour | 5 | 72.4 | Vertex peak amplitude left sensory peak amplitude, vertex peak latency |

